# Effective excitability captures network dynamics across development and phenotypes

**DOI:** 10.1101/2024.08.21.608974

**Authors:** Oleg Vinogradov, Emmanouil Giannakakis, Victor Buendía, Betül Uysal, Shlomo Ron, Eyal Weinreb, Niklas Schwarz, Holger Lerche, Elisha Moses, Anna Levina

## Abstract

Neuronal cultures *in vitro* are a versatile system for studying the fundamental properties of individual neurons and neuronal networks. Recently, this approach has gained attention as a precision medicine tool. Mature neuronal cultures *in vitro* exhibit synchronized collective dynamics called network bursting. If analyzed appropriately, this activity could offer insights into the network’s properties, such as its composition, topology, and developmental and pathological processes. A promising method for investigating the collective dynamics of neuronal networks is to map them onto simplified dynamical systems. This approach allows the study of dynamical regimes and the characteristics of the parameters that lead to data-consistent activity. We designed a simple biophysically inspired dynamical system and used Bayesian inference to fit it to a large number of recordings of *in vitro* population activity. Even with a small number of parameters, the model showed strong inter-parameter dependencies leading to invariant bursting dynamics for many parameter combinations. We further validated this observation in our analytical solution. We found that *in vitro* bursting can be well characterized by each of three dynamical regimes: oscillatory, bistable, and excitable. The probability of finding a data-consistent match in a particular regime changes with network composition and development. The more informative way to describe the *in vitro* network bursting is the effective excitability, which we analytically show to be related to the parameter-invariance of the model’s dynamics. We establish that the effective excitability can be estimated directly from the experimentally recorded data. Finally, we demonstrate that effective excitability reliably detects the differences between cultures of cortical, hippocampal, and human pluripotent stem cell-derived neurons, allowing us to map their developmental trajectories. Our results open a new avenue for the model-based description of *in vitro* network phenotypes emerging across different experimental conditions.

## INTRODUCTION

Networks of neurons *in vitro* are a minimal system that enables the study of the interaction between different levels of neuronal network organization [1, 2]. As an experimental model, cultures of neurons have been used to identify mechanisms of neuronal plasticity [3, 4], homeostatic adaptation [5], E*/*I balance [6–8], network development [6], and principles of collective activity [9, 10]. More recently, with the development of stem cell technologies, networks of human pluripotent stem cell (hPSC)-derived neurons *in vitro* allowed researchers to directly study how disease-associated genotypes affect the properties of single neurons and the networks they form [11–13].

Mature networks of dissociated neurons *in vitro* robustly exhibit coordinated population bursting activity [14–17]. This activity usually manifests as large synchronous events propagating through the whole network, followed by long, irregular periods of quiescence. Such network activity has been shown to occur in cortical, hippocampal, striatal, and spinal cord cultures *in vitro* as well as cultures of both induced and embryonic hPSC-derived neurons [18–22]. Population bursting activity is one of the markers of successful network development [23] and is often used to evaluate the effects of experimental conditions on neurons and networks [13, 21, 22, 24–28].

Population bursting can be modeled as a low-dimensional slow-fast dynamical system [16, 29, 30]. Theoretical studies suggest that network bursting emerges from an interaction of fast recurrence and an adaptation mechanism that slowly adjusts network excitability [9, 16, 20, 24, 29, 31]. This principle can be described as a low-dimensional approximation of the recurrent activity driven by noise [32, 33].

Fitting a low-dimensional dynamical system to recorded network dynamics can provide insights into the mechanisms of population dynamics and can help uncover the differences between experimental conditions. Several studies showed that fitting reduced models allows us to map out the patterns of cortical network activity *in vivo* and *ex vivo* and suggests mechanistic explanations of the transition between the activity regimes induced by anesthesia, sleep and wakefulness [33– 35]. This approach has several benefits. The data-fitted model parameters can be rendered more interpretable. Additionally, it allows us to analyze the data using dynamical systems theory, map dynamical regimes of network activity, and identify the phase transitions. However, major challenges in finding interpretable and appropriately describing the data parameters prevented this approach from being systematically applied to the recordings of network activity. First, finding data-consistent parameters for such models for large volumes of data remains a daunting task. Second, when using expressive models, many parameter combinations describe the data, and thus, finding an effective parametrization that allows the interpretation and leveraging of this redundancy becomes a necessity. In this study, we show how both these challenges can be solved to generate interpretable models that allow us to analyze the differences between culture types and network development.

Here, we establish a simplified population model for network bursting activity *in vitro*. We show that the *in vitro*-like bursting activity emerges in a range of dynamical regimes and can be governed by excitable, oscillatory, or bistable dynamics. We applied simulation-based inference [36] to approximate the distribution of model parameters that lead to data-consistent model dynamics. This allowed us to access the dependencies between model parameters and helped us establish an effective excitability metric, that summarized the level of intrinsic drive and adaptation in the network. We fitted the model to a wide range of recordings of rodent primary cortical and hippocampal cultures as well as cultures of hPSC-derived neurons. Although they all display very similar network bursting activity, we show differences in their dynamical regimes and effective excitability both in the mature state and during development. Finally, we demonstrate that changes in the effective excitability can be induced by acute perturbations of single cell excitability *in vitro*. Our results establish a way to describe the population activity of cultures of neurons *in vitro* using low-dimensional, data-consistent models that directly map parameters to observed dynamics, thereby enabling the effective representation and discovery of changes across various experimental conditions and phenotypes.

## RESULTS

### A low-dimensional rate model with slow adaptation generates bursting activity

Network activity in living neuronal networks *in vitro* typically consists of synchronous events that rapidly propagate throughout the whole network and long inter-burst intervals of irregular firing with low firing rate. We model this activity using a simplified phenomenological model that allows us to pin down the dynamical regimes associated with bursting activity and define the key parameters controlling the activity (Fig. 1). The average network activity (*x*(*t*)) and the adaptation current (*w*(*t*)) follow:

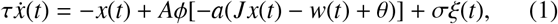

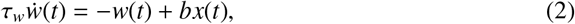

where *ϕ* is the sigmoid nonlinearity *ϕ*(*z*) = (1 + *e*^*z*^)^−1^, ξ(*t*) is an uncorrelated Gaussian white noise, and *σ* defines the noise intensity. Here, τ sets the fast neural timescale, *A* is the scale of the nonlinearity, *a* defines the gain of the input-output relationship, *J* is the strength of recurrent interaction, and *θ* sets an intrinsic drive (for cultures can be interpreted as level of spontaneous activation). We interpret the noise in the model, *σξ*, as fluctuations occurring within the recurrent population of neurons, e.g., due to the spontaneous firing of neurons. The adaptation current accounts for the average neuron’s adaptation [37, 38], *τ*_*w*_ sets its timescale, and *b* controls the adaptation strength. The population firing rate is computed as *y*(*t*) = exp (*mx*(*t*)) where *m* is found via Poisson regression (see Methods I A).

**FIG. 1.**
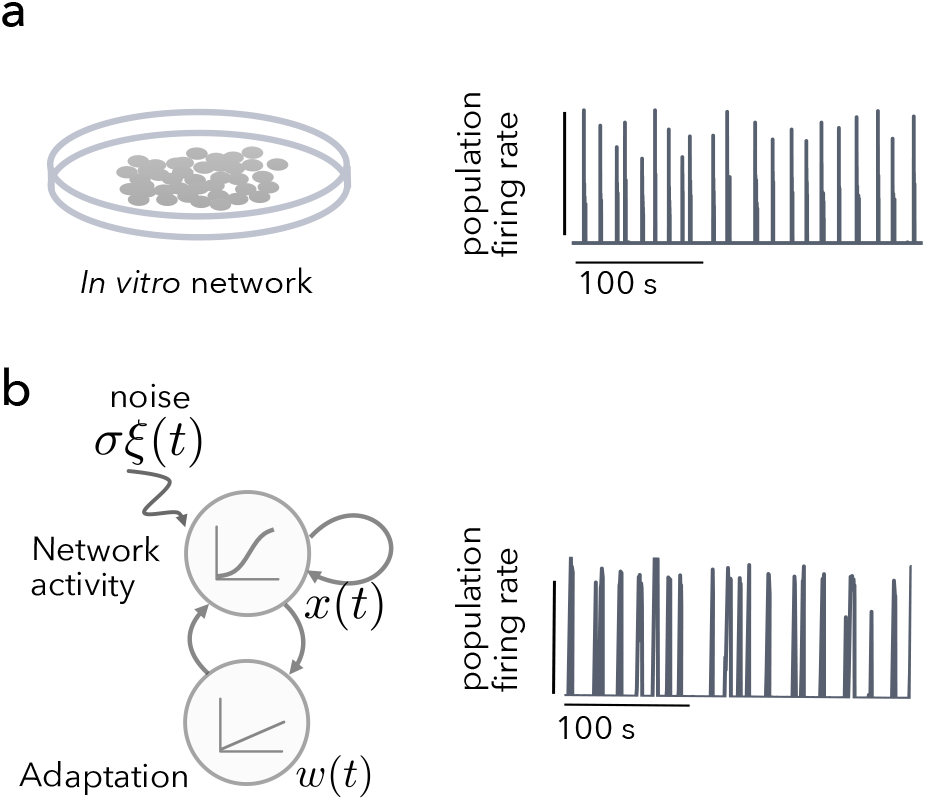
Recurrent rate model with slow adaptation produces *in-vitro-* like population bursting dynamics **a)** Mature cultures of dissociated primary mouse neurons robustly exhibit population bursting *in vitro* (firing rate is averaged over 64 channels in 100ms bins, data are from [25], cortical culture 21 DIV). **b)** Schematic of the recurrent rate model with slow adaptation driven by noise and an example of the population activity produced by the model.

The model can reproduce the temporal statistics of network bursting activity emerging in a population of neurons. Varying four key parameters – the intrinsic drive *θ*, the adaptation strength *b*, the adaptation timescale τ_*w*_, and the noise intensity *σ*, allows us to find bursting-like activity matching the main summary statics of network bursting in the data. Specifically, we are looking for the parameters that match mean *inter-burst intervals (IBI)*, variability of inter-burst intervals measured as the *coefficient of variation* (CV) of IBI, and average *burst duration*.

### Network bursting activity emerges in noise-driven excitable, bistable, and oscillatory regimes

The minimal model can be well-characterized in terms of the nature and number of fixed points. Depending on parameters, it presents a limited set of dynamical regimes [39], some of which produce network bursting-like dynamics.

Among the four key parameters of our model, only the excitability and adaptation strength control the number and stability of fixed points, and we use them as primary parameters to visualize the bifurcation diagram (Fig. 2a). The system has three types of solutions supporting bursting-like activity: an oscillatory solution manifesting as a limit cycle, an excitable solution with one stable fixed point (corresponding to high or low activity), or a bistable solution with three fixed points (two stable and one unstable in-between) (Fig. 2b and see Methods I D).

**FIG. 2.**
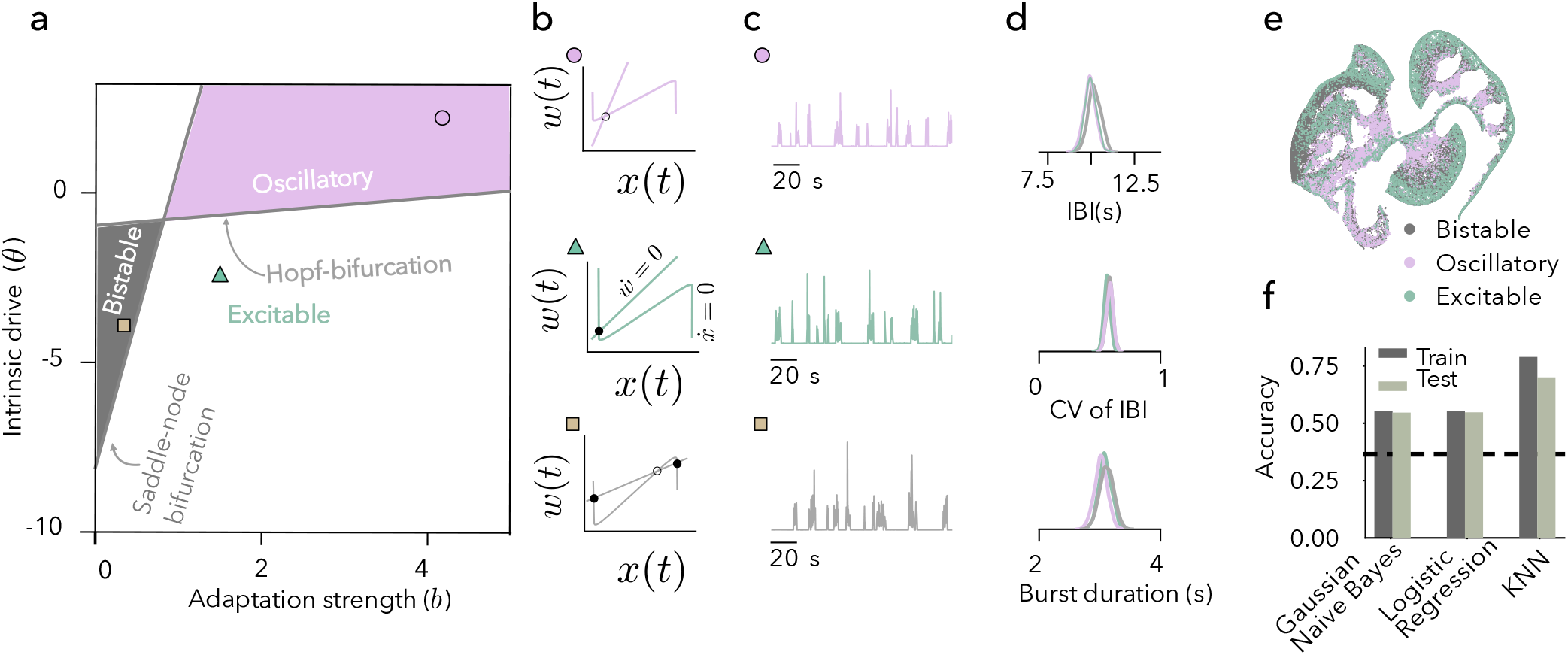
Multiple dynamical regimes produce similar network bursting activity. **a)** The bifurcation diagram of the model for changing the intrinsic drive (θ) and adaptation strength (*b*). **b)** Nullclines of the model in the oscillatory, excitable, and bistable regimes **c)** Examples of generated spike counts for the model in three dynamical regimes fine-tuned to produce similar activity. **d)** Distributions of inter-burst intervals (IBI), CV of IBI, and burst durations for the three example sets of model parameters (200 simulations with different random seeds). **e)** t-SNE embedding of the bursting summary statistics (inter-burst intervals, CV of inter-burst intervals, and burst duration) color-coded according to the dynamical regimes does not show any clear boundary between the regimes. **f)** Classification performance increases in classifiers that rely on local similarity of points (K-Nearest Neighbors) rather than convex boundary (Linear Regression and Gaussian Naive Bayes) between the regimes.

All three types of solutions can generate a network-bursting activity with a wide range of average inter-burst intervals, CV of inter-burst intervals, and average burst durations (See Extended Figures Fig. 10). Bursts in these three cases are generated by different mechanisms. In an oscillatory regime, the system deterministically oscillates between the down and up states, the latter corresponding to the network burst. A strong noise can perturb such oscillations and make them irregular (Fig. 2b and c, pink, 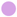). In the case of a single fixed point, network bursting can occur as an instance of excitable dynamics: the noise drives the model away from the fixed point in a direction that requires passing a high activity region (that corresponds to the burst) before slowly returning to the fixed point (Fig. 2b and c green, 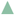). Lastly, bursting can emerge as noise-driven transitions between two stable fixed points (one of which corresponds to the higher activity and bursting) in the bistable regime (Fig. 2b and c, gray, 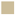).

Points in the parameter space belonging to different dynamical regimes can show very similar activity and corresponding summary statistics (Fig. 2d right). To characterize this similarity, we use the summary statistics of the model’s activity (inter-burst intervals, CV of inter-burst intervals, burst duration, for details see burst extraction methods in Methods I A) to determine which regime gave rise to this statistics with unsupervised and supervised methods. A visualization of the dynamical regimes with t-SNE (Fig. 2e) indicates that regimes are well-concentrated but without clear convex and continuous boundaries. Both linear (Logistic Regression) and nonlinear (Gaussian Naive-Bayes) classifiers perform only slightly better than chance. The KNN-based classifier uses only local properties of individual observations and thus can reach better performance; however, it is still below 75% accuracy on the test set (Fig. 2f).

### Invariant bursting dynamics across different dynamical regimes

We showed in the previous section that it is impossible to establish a one-to-one correspondence between the summary statistics and model parameters. Additionally, when fitting models to the experimental observations, the summary statistics are noisy due to the limited data availability. Thus, instead of searching for a single best parameter set for each given burst statistic, we aim to determine the whole distribution of parameters that could lead to the observed data called *a posterior distribution*. To approximate the posterior distribution, we employ a machine learning technique called simulation-based inference [40]. The idea is to simulate a model with many different parameter sets and gather the summary statistics of model observables. We are interested in inverting this map so that we get the distribution of model parameters that could lead to each combination of summary statistics. In most cases, this inversion cannot be done analytically, and the machine learning community is spending a lot of effort optimizing the numerical procedures and number of simulations needed for reliable inversion [41]. Here, we use an amortized inference approach and train a neural network to approximate the inversion [40]. The fitted posterior can be evaluated for any given set of experimentally observed summary statistics (in our case, mean inter-burst intervals, CV of inter-burst intervals, and burst durations) and return the model parameters that produce the activity with these features.

We estimated the posterior distribution of parameters given the summary statistics of network bursting in cortical cultures we recorded at 17 and 18 DIV using MEA (example recording seen in Fig. 3 a top, for recording details see Methods II). The fitted neural posterior reliably identified the model parameters that produce the desired bursting summary statistics (Fig. 2 a, traces 1 and 2). We illustrate this for the experimental data sample by simulating models with parameters drawn from the posterior and observe that the simulated statistics are narrowly centered around the experimentally observed values (Fig. 3 b), and deviating from the posterior distribution leads to the large mismatch in the produced data (Fig. 3 a, trace 3, Fig. 3 b red marks).

**FIG. 3.**
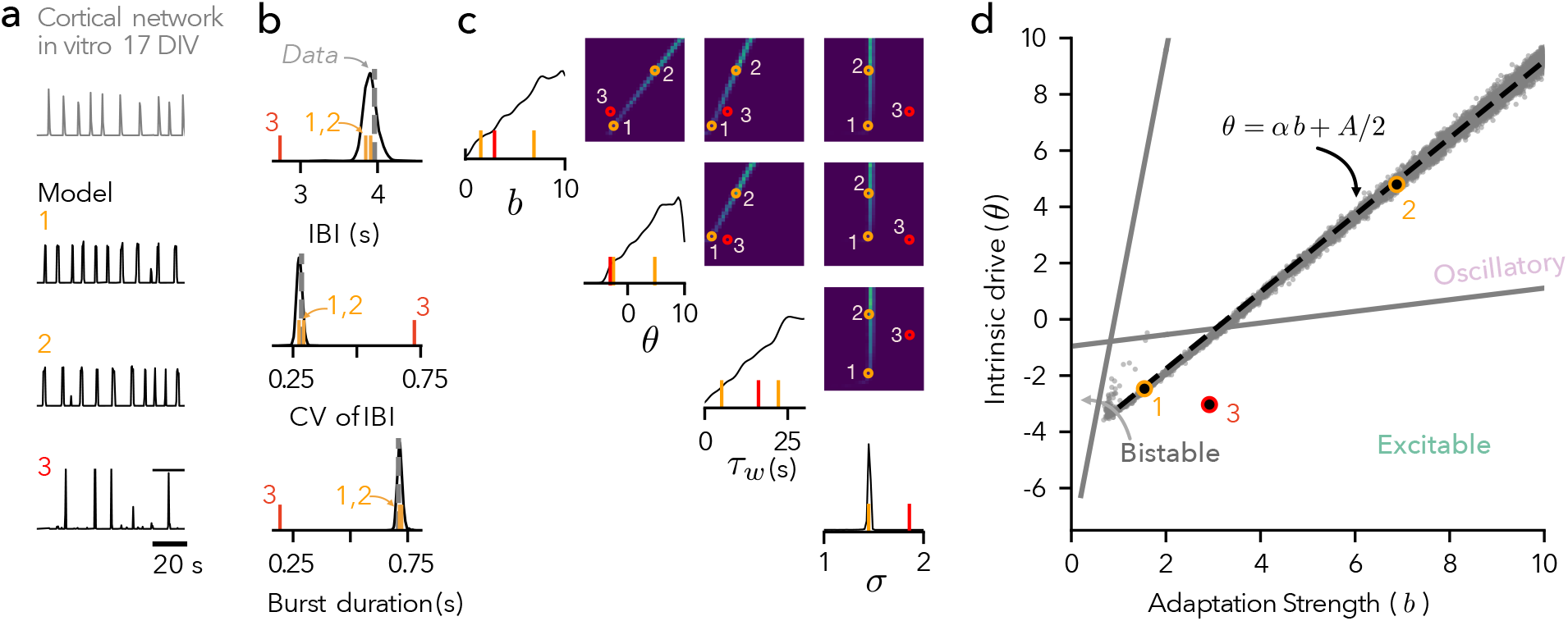
Simulation-based inference of the data-consistent model parameters identifies the model invariances and suggests a simple effective excitability metric. **a)** Example of firing rate averaged over recording channels from a mouse primary cortical culture at 17 DIV on MEA and examples of model activity with the parameters sampled from the approximated posterior given a set of summary statistics (1,2-yellow) and outside of it (3 - red). **b)** Summary statistics of bursting activity from the posterior (black lines: histograms of burst statistics generated by 2000 models with parameters drawn from posterior distribution) closely match the summary statistics of bursting activity from the data (vertical dashed line). The example points from **a** are marked by corresponding colors and legends. **c)** Pairwise plots of samples from the approximated posterior distribution for bursting in an example cortical culture (17 DIV) obtained with simulation-based inference (see main text) show a linear dependency between the intrinsic drive and adaptation. **d)** Dependency between the intrinsic drive and adaptation strength on top of the bifurcation diagram. The dashed line shows an analytically derived dependency line. The slope of this line (*α*) defines the effective excitability.

The approximated posterior distribution of model parameters revealed the dependencies between the parameters that allow for invariant dynamics across different underlying dynamical regimes. To identify them, we can look at the pairwise marginal distributions, where the dependencies between parameters are visible in the specific, anisotropic density (Fig. 3 c). The posterior distribution showed an almost linear dependency between the network excitability and the adaptation parameter (Fig. 3 d). The system can maintain the same bursting features when excitability increases by strengthening and slowing the adaptation current. Thus, for varying values of network excitability, one can find corresponding adaptation parameters that would maintain invariant bursting activity. For this example of summary statistics, the posterior distribution spans across different dynamical regimes (mainly oscillatory but also excitable).

### The statistics of network bursting determines the effective excitability in a fitted model

In the previous section, we observed that a continuum of models can generate a given network bursting activity, which is possible due to the striking dependencies between the model parameters. To explain this dependency analytically, we further simplified the model by substituting the input-output nonlinearity in Eq. 2 with a piece-wise linear approximation and assumed a separation of timescales *τ* ≪ *τ*_*w*_. This allowed us to describe analytically the network dynamics for given *θ* and *b* and obtain the mean burst duration and mean inter-burst interval. In brief, we can find the values of network activity and adaptation at the start/end of the burst as a local minimum/maximum of the 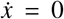 nullcline. Given these values, we can obtain exactly the decay of the adaptation variable after the beginning of an inter-burst interval (that determines the length of this interval). Analogously, we analyse the rise of *w*(*t*) to obtain the burst duration. Inverting this solution, we find the relationship between *θ* and *b* for each mean interburst interval and burst duration. Then, by combining these equations, we find a deterministic solution that describes the relationship between *θ* and *b* in a full model with noise and without timescale separation (see dashed line in Fig. 3 d). For more details on the analytics see MethodsI E.

Our analytic solution describing the relationship between excitability and adaptation strength for relevant parameter ranges can be closely approximated by the line

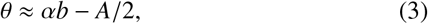

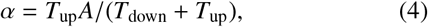

where *T*_up_ is the mean burst duration and *T*_down_ is the mean inter-burst interval (See Methods I E). We define the slope of this line (*α*) as the effective excitability (Fig. 3 d, dashed line), which can be directly computed from the summary statistics of network bursting. This value closely follows the slope of the dependency between the intrinsic drive and adaptation obtained with simulation-based inference (Extended Figures Fig.7).

### The effective excitability captures the differences between cortical, hippocampal, and human PSC cultures

Next, we asked if the differences in bursting dynamics between different types of cultures are reflected in the parameters of the fitted models. We analyzed the model parameters that reproduce the bursting statistics for primary mouse cortical and hippocampal cultures [25], and hPSC-derived neuronal cultures [21]. We first focused on the bursting activity of mature cultures (older than 14 DIV for mice cultures [25] and after 21 days for hPSC [21]).

The parameters of the fitted models are clearly different (practically non-overlapping) between different types of cultures (Fig. 4a) and can capture well the activity of the respective cultures (Fig. 3b). One of the salient differences is related to the preferred type of the dynamical regime. We estimated the fraction of parameter samples for each culture type that fall into bistable, excitable, and oscillatory regimes. We found most of the samples for developed cortical cultures are most consistent with the oscillatory dynamics. The hippocampal and the hPSC cultures are primarily found in the excitable regime.

**FIG. 4.**
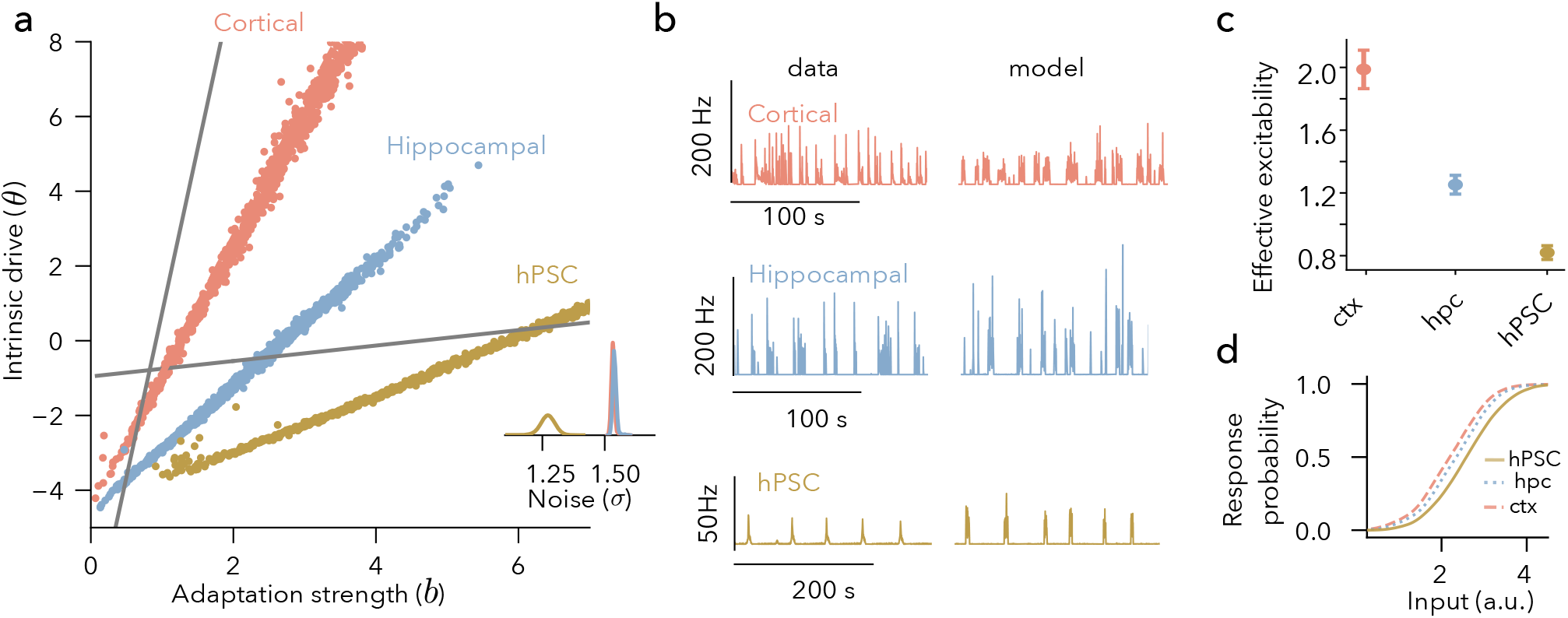
The effective excitability captures the differences between primary mouse cortical and hippocampal cultures as well as hPSC-derived neuron cultures. **a)** Samples from posterior distributions of the model parameters fitted to example cortical and hippocampal cultures at 24 DIV [25], and hPSC cultures at 38 days on MEA [21, 42]. **b)** Average firing rate of the example cultures and the corresponding model activity (fitting rate is computed in 100ms time bins). **c)** Effective excitability in cortical, hippocampal culture (older than 14 DIV), and hPSC cultures older than 21 Days on MEA (error bars - s.e.m., p < 0.00001, pairwise ind. t-test cortical vs hippocampal cultures t-val = 8.8, p < 0.00001; cortical vs hPSC t-val = 9.158 p < 0.00001; hippocampal vs hPSC t-val=8, p < 0.00001). **d)** The differences in the effective excitability predict different network response properties in cortical, hippocampal, and hPSC cultures. Example response curves computed for the cultures from **a, b**.

As previously shown, the linear dependency accurately describes the marginal posterior distribution of θ and *b*, and thus, we can use the effective excitability (*α*, Eq. 4) to capture the essence of the different model-fits. Considering all mature recordings in the datasets (n=150 for cortex, n=221 for hippocampus, and n=92 for human PSC), we see a clear statistical difference between them: in hPSC cultures, the effective excitability is significantly lower than in hippocampal cultures that, in turn, is lower than in cortical cultures (Fig. 4c). We also found systematic differences in the noise level between hippocampal, cortical, and, hhPSC cultures, but not in the timescale of adaptation (see Extended Figures Fig. 12).

We verify that the effective excitability controls the network’s responsiveness to external stimulation by applying short inputs of increasing strengths. We perturbed the system by applying an external stimulus at different times within the inter-burst intervals and finding the probability of such input to initiate a burst that we called response probability. Changing the effective excitability shifted the response curves to the left, thus increasing the model’s responsiveness (Extended Figures Fig.. 15 and see also Fig. 4d). We computed the response probabilities for the models fitted to the hippocampal, cortical, and hPSC cultures (Fig. 4d). As predicted by the effective excitability, the responsiveness was the highest in cortical cultures and the lowest in hPSC cultures (Fig. 4d).

### Mapping the developmental trajectory of the bursting activity with models parameters

Bursting dynamics is known to strongly change throughout development, reflecting neuronal maturation, synaptogenesis, and myelination of axons [6, 19, 21]. Maturation of a network is associated with increasing network excitability measured as a decrease in the stimulation threshold needed to elicit a burst [6, 14]. We asked if the effective excitability ratio estimated from the spontaneous activity of networks throughout the development would reflect these changes. We further analyzed the data for cortical, hippocampal, and hPSC cultures, now considering all recording days. [25, 42]. For the recordings of cortical and hippocampal cultures, the network activity was detectable from approximately 7 DIV and was then recorded until 28 DIV. We could identify the first bursting activity in hPSC cultures from around 18 days after plating on MEA.

The effective excitability estimated from the model fitted to spontaneous activity of standard mouse cortical and hippocampal cultures showed a smooth increase over the course of development. The activity of cortical and hippocampal cultures first emerges as sparse and highly irregular, with large inter-burst intervals and small burst durations. This is reflected in small values of effective excitability (Fig. 5a). The models that fit the data are then predominately found in the excitable region (Fig. 5a inset). Throughout development, the activity increases, bursts become more frequent, and their duration increases, which is reflected by an increase in the effective excitability (Fig. 5). The parameters of the model accordingly move closer to a Hopf-bifurcation, and at the later stages, some of the networks are predominantly consistent with oscillatory dynamics.

**FIG. 5.**
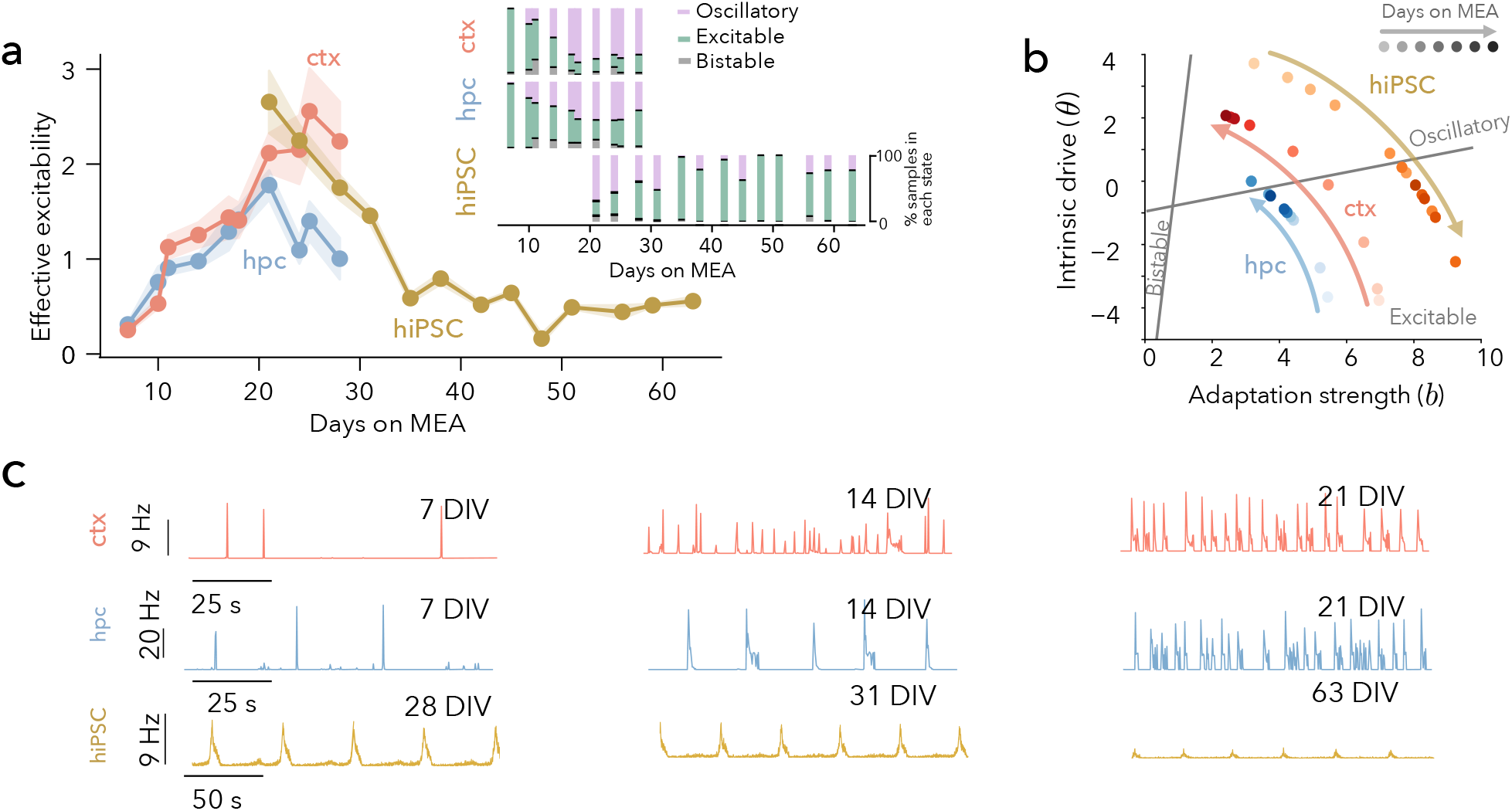
The effective excitability allows us to trace the developmental differences in the collective bursting dynamics between different preparations. **a)** Both primary mouse cortical and hippocampal cultures [25] show an increase in effective excitability. In contrast, the cultures of hPSC-derived neurons show a gradual decrease in network excitability followed by a long plateau [21]. **Inset** shows the fraction of samples from the posterior distribution in each regime per day and culture type. **b)** Sketch of the changes in the effective excitability for the three culture types. The models suggest that during network development, hippocampal and cortical cultures may undergo a phase transition from the excitable to the oscillatory regime; hPSCs that we analyzed first exhibit network bursting that is consistent with the oscillatory regime and then move towards excitable dynamics. **c)** Examples of average firing rates (in 100ms time bins) at different times during development.

When bursting first appears at 7 DIV, cortical and hippocampal cultures are not distinguishable in terms of their excitability. Around 20 DIV the cortical and hippocampal cultures diverged in their development with the cortical cultures, on average, showing a higher effective excitability (Fig. 5).

The cultures of hPSC-derived neurons showed a different pattern of development. We could reliably detect network bursting activity only at 18 DIV. The network activity at this point was characterized by a large effective excitability (Fig. 5). Then, over development, hPSC networks slowly decrease the ratio and reach a plateau after 31 DIV. Thus, the development of collective activity follows a different trajectory compared to the primary neuron cultures. However, our model suggests that the hPSC-derived neuronal cultures still exhibit either oscillatory or excitable dynamics, and at different stages of their development, it could be consistent with primary cultures of dissociated neurons.

### Acute perturbations of KCl concentration *in vitro* are captured by the changes in effective excitability

Finally, we examined how the inferred effective excitability is related to the acute changes in excitability of individual neurons. We cultured rat primary hippocampal neurons and recorded their activity in different concentrations of KCl that is known to change the excitability of neurons [18, 43]. We first recorded the activity in standard, close to physiological, 4mM KCl. Then we changed the whole medium to a new concentration: 1, 2.5, 5.5, 7.0, 10.0 mM (See Methodsfor experimental details).

The fitted models’ effective excitability mirrored expected changes in excitability from varying KCl concentrations. When increasing the concentration of KCl between 1–5.5mM the model effective excitability increased and saturated at 7mM (Fig. 6a). When the concentration was further increased to 10mM the networks typically stopped showing collective bursting (Extended Figure 16). Thus, we show that the estimated effective excitability can effectively read out the changes introduced by acute perturbations of single neuron excitability reflected in the changes of network activity.

**FIG. 6.**
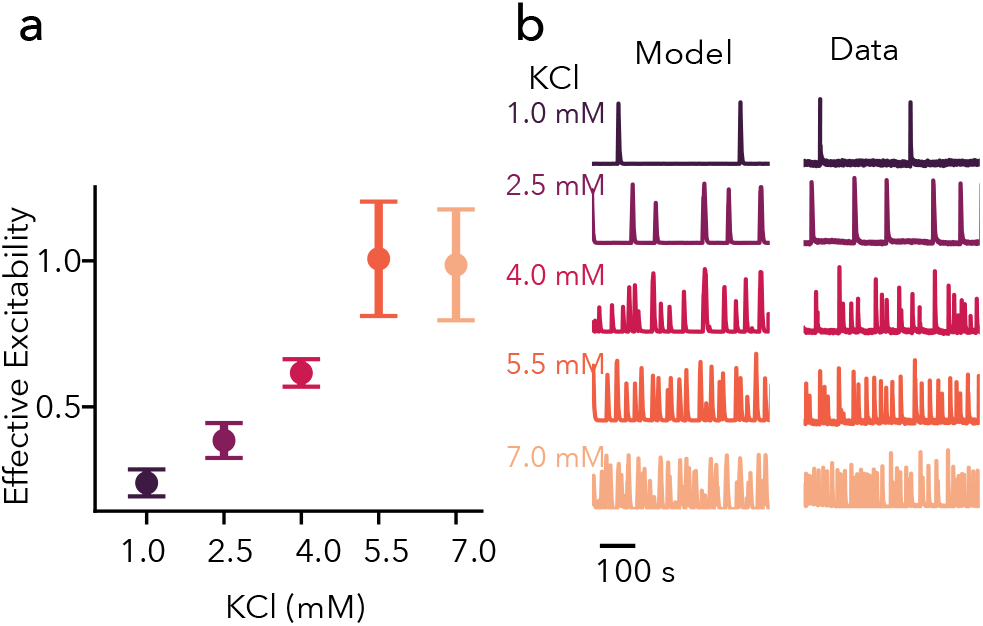
The measured effective excitability reflects the changes in neuronal excitability induced by an acute manipulation of the KCl concentration. **a)** Average effective excitability of hippocampal cultures at different concentrations of KCl in the medium (error bars - s.e.m.). Compared to the recordings at 4mM KCl, the estimated effective excitability was significantly higher when the KCl was increased (5.5, 7mM t-test with 50000 permutations p=0.012 and 0.012 with BH-FDR correction). The estimated effective excitability was significantly lower in 1 and 2.5mM of KCl (p=0.0079, 0.038 with BH-FDR correction). We did not see significant differences between 5.5 and 7mM KCl. **b)** Example of the activity of the culture in increasing concentrations of KCl with the corresponding best-fitting models.

## DISCUSSION

We demonstrated that the effective excitability, defined as the ratio of the network drive and adaptation is one of the key determinants of network bursting behavior. To this end, we used a simplified dynamical systems model and a combination of Bayesian inference and analytic techniques. This effective excitability separates different types of cell cultures and captures the developmental differences in population activity. We show analytically and confirm numerically that as long as the effective excitability remains the same, the model exhibits invariant bursting behavior in dynamical regimes with diverse underlying parameters. Thus, we identified the essential invariance that allows us to capture the most important features of the bursting dynamics with a minimal number of parameters.

The excitable, oscillatory, and bistable network dynamics that we identify as main possible regimes of network bursting have been previously studied in the context of cortical and hippocampal *in-* and *ex vivo* activity such as up-down states, slow-wave oscillations and sharp wave ripples [32, 33, 35]. Our results are closely related to the work of Levenstein et al. 2019 [33], which focused on the dynamical regimes of the cortex and hippocampus during NREM sleep *in vivo*. By matching the distributions of durations of up and down states they demonstrated that mesoscopic activity of cortex and hippocampus during NREM is best explained by excitable dynamics and suggested a mechanism of sharp-wave ripple-slow wave coupling. Our model is similar to the adapting recurrent activity model they propose, with comparable dynamical regimes and bifurcations. However, instead of multiplicative noise with autocorrelations, we use additive Gaussian white noise. Additionally, we only consider a linear adaptation that enabled us to explicitly find the relationship between the network bursting activity and model parameters. In our paper, we extend their results by fitting the four key parameters of the model simultaneously. This allows us to identify parameter dependencies that allow for statistically identical dynamics in different dynamical regimes beyond the excitable state. This result further suggests that there are multiple paths to network bursting dynamics, which can be critical in context of interpreting the large variability observed in cultured neuronal networks [14] and unifies varies mechanisms suggested to explain network bursting *in vitro* [9, 16, 29, 44].

A detailed analysis of the dependencies between the model parameters helped us to identify the effective excitability parameter. This parameter can be directly calculated from the network burst durations and inter-burst intervals. We show that the effective excitability discriminates well different types of cultures and captures their developmental profiles. For example, we found differences between the effective excitability of both mature and developing cortical, hippocampal, and hPSC cultures using publicly available datasets [21, 25]. Our results are consistent with the original publications, where differences were detected using individual bursting summary statistics and dimensionality reduction of a large vector of bursting features. In contrast, our model-based effective excitability offers a more compact and interpretable description of the activity and can be directly used to predict the responsiveness to external stimulation. Importantly, the differences in responsiveness to external stimulation between cortical and hippocampal cultures that we identify using the model were indeed shown in experiments [6, 14, 45]. The response properties of hPSC cultures remain to be shown. Interestingly, compared to primary cultures of hippocampal and cortical neurons, hPSC cultures reach their steady state of effective excitability via a different developmental trajectory: they start at high levels of excitability and most likely exhibit early bursting driven by an oscillation. Matching the developmental trajectories of hPSC to identify the differences with primary cultures of neurons is one of the challenges in the field of hPSC-based disease models [21, 46]. Future studies should focus on comparing the developmental profiles between various protocols for hPSC-derived neurons. Here, our approach can help align the staging for different preparations and protocols using the effective excitability.

In our work, we focused on a simplified modeling approach to uncover the principles that constitute the dynamics [33, 35, 47–50]. A complementary approach is to use very detailed models, which can potentially reflect more detailed features of the data and can also be parameterized using experimentally tractable parameters[16, 24, 51]. However the the costs of finding the proper parameter set in this very highdimensional set may be prohibitively high: to capture the dynamics of a single cortical neuron one might need thousands of equations [52] or thousands to millions of parameters in machine learning models [53, 54]. Additionally, it remains to be determined how to use the invariances found in a very highdimensional space to generate verifiable predictions. Merging the simplified and extended approaches will be a very promising avenue for future research.

Due to its simplicity our modeling ignores some aspects of network bursting variability such as burst shapes [14], and compositions of busts [55]. The model also does not explicitly include inhibition, assuming that inhibitory interaction happens on a very short timescale and is captured by the variance of the noise [7, 24]. Furthermore, our model does not include the notion of space and is therefore not suitable for describing the spatial phenomena in network dynamics [9, 27].

Our work demonstrates that simplified models are not only useful for describing the dynamical regimes underlying network activity but also for building a deeper quantitative understanding of the statistics of network activity. Furthermore, we show that in the context of *in vitro* networks, mapping their population activity to model parameters allows for a rapid assessment and interpretation of the emerging network phenotypes.

## ACKNOWLEDGMENTS

This work was supported by a Sofja Kovalevskaja Award from the Alexander von Humboldt Foundation. OV thanks the International Max Planck Research School for the Mechanisms of Mental Function and Dysfunction, the Joachim Herz Foundation, and Feinberg Graduate School for their support. EG thanks the International Max Planck Research School for Intelligent Systems (IMPRS-IS) and the Tübingen AI center for their support. We acknowledge the support from the BMBF through the Tübingen AI Center (FKZ: 01IS18039B). AL is a member of the Machine Learning Cluster of Excellence, EXC number 2064/1 – Project number 39072764. The authors thank Tim Schäfer, Roxana Zeraati, Richard Gao, Michael Deistler, Jakob Macke, Paul Spitzner, Tanguy Fardet, Johannes Zierenberg, and Dominik Freche for insightful discussions.

## I. METHODS

### A. Rate model

The rate model (Eq. 2) was simulated using the Euler-Maruyama method with a 0.05 ms integration time. To extract the summary statistics, we first simulate 10s of activity and exclude this as a burning-in period. Then we keep simulating until at least 30 bursts are collected.

#### Burst detection

The bursts were detected by discretizing the activity values *x*(*t*) and generating quasi-spike times where *x* > 0, with the number of spikes at that time being equal to the discretized *x*. We use a modified version of the minimum interval algorithm (MI), which has been shown to robustly detect burst [24, 56]. The algorithm detects bursts as the activity during which inter-spike intervals are below a threshold. Specifically, we set the inter-spike interval threshold to be 10ms and check if the bursts are longer than 10ms and if the number of spikes within the bursts is larger than 5. We then merge the bursts if the inter-burst interval between them is longer than 20ms.

#### Burst summary statistics

From the detected burst times, we compute the average burst durations (*T*_up_) and the duration of inter-burst intervals (IBI, *T*_down_) as the time between the end of a burst and the beginning of the next one (note that we use a similar definition for the burst analysis of experimental data). We also estimate the coefficient of variation of the inter-burst intervals (as std(*T*_down_)/mean(*T*_down_)).

#### Spike rate mapping

After simulating the activity of the model, we mapped on it the neuronal firing rate as *y*(*t*) = exp (*mx*(*t*)), which is inspired by the idea of cascade models [57] and the single neuron spike response model [58]. We fit the parameter *m* directly from experimental data with Poisson regression using the average spike counts within each burst. The rate of the inhomogeneous Poisson process is controlled by the average activity *x*(*t*) and *m* scales it to match the spike counts in the recording.

#### Input-response of the model

To numerically analyze the model responses to external inputs (Fig. 4d), we numerically simulated the model for 6000s and discarded the first half of the simulation. We then randomly choose 200 points with different x(t) and w(t) from the down-state (e.g. between bursts) values and treat them as independent initial conditions. We run 200 independent simulations with these selected initial conditions with increasing external input (from 0 to 6). The input is applied within nonlinearity, so the original equation (Eq. 2) is modified as ϕ[−*a*(*Jx*(*t*) − *w*(*t*) + θ + *μ*)], where *μ* is external input. The stimulation is applied to 10ms after which we check if it evoked a burst. We repeat this process 100 times with different random seeds for the external white noise.

### B. State classification with summary statistics

We tested if we could identify the dynamic state of the model from the bursting summary statistics. We sampled 150000 model parameters in the wide range (*b* [0;10], θ [-10;10], log(*τ*_*w*_)[0;14s], *σ* [0;2]) and fit a set of linear classifiers (Linear Regression and Gaussian Naive Bayes) as well as KNN-based classifier to predict the state based on the IBI, CV of IBI, and burst duration (Fig. 1f). The classifiers were trained on 80% of the data and tested the remaining 20%.

### C. Simulation-based inference of the model parameters

#### Conditional density estimation

To directly determine how the bursting statistics map onto the model parameters, we use a simulation-based inference approach. We use the simulations to build a conditional density estimator p(parameters|summary) that can be evaluated for other summary statistics without additional sampling [40, 41].

As a conditional density estimator we use a neuronal spline flow [59] within the SBI-toolbox [60]. The model included 5 transformations and 50 features per layer.

We train the conditional density estimator using 300,000 samples from a wide prior of parameters (Table I). The estimator allows us to obtain an approximate posterior probability of the model parameters given a set of summary statistics (p(parameters|IBI, CV of IBI, duration)). We evaluated the predictive performance of the model and found that it achieves excellent results for a wide range of summary statistics (Fig. 11). We run the predictive checks analysis by evaluating the posterior probability of parameters for the experimental data, simulating the model with these parameters and comparing the resulting values (Fig. 11).

**TABLE I.** Prior values for parameters. The values of τ_*w*_ were sampled in the log space *log*(200) to *log*(200000)

| Parameter | Lower Bound | Upper Bound |
| --- | --- | --- |
| $b$ | 0.05 | 20.0 |
| $\theta$ | -10.0 | 15.0 |
| $\tau_w$ | 200.0 ms | 200000 ms |
| $\sigma$ | 0.01 | 2.0 |

#### Fraction of states per observation

We compute the changes in the fraction of states consistent with the experimental data (Fig. 5 inset). For each summary statistic of the experimental observation, we randomly sample 10,000 parameters from a posterior distribution for each observation, pool these samples together, and calculate the fraction of parameters that fall into each state (bistable, excitable, and oscillatory).

### D. Stability analysis of the rate model

We study the phase portrait of the model, using standard stability analysis for the deterministic case [39]. The system is given by

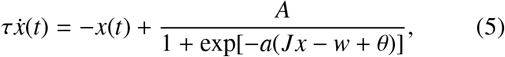

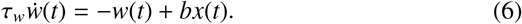

We analyze the case where *τ* = 1 and *J* = 1. We start by computing the fixed point and nullclines of the model. The nullclines are given by

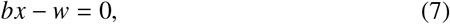

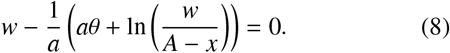

The fixed points *x*^*^, *w*^*^ are given by the intersection of two lines and can only be estimated numerically. The solutions are typically 1 or 3 fixed points (Fig.1c). The stability of the fixed points can be calculated by linearizing the system around the fixed points:

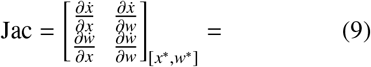

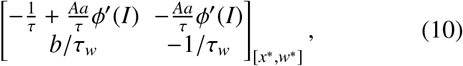

where the derivative of the sigmoid is 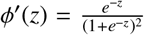 and *I* = −*a*(*θ* − *w* + *x*)). The signs of the eigenvalues of the Jacobian determine the stability of fixed points,

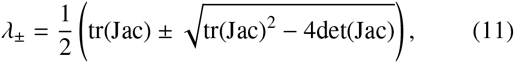

where tr is a matrix trace and det stands for the matrix determinant. The determinant of the matrix is given by

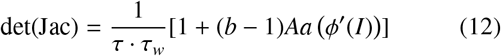

and the trace reads

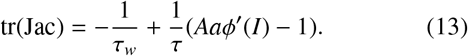

By analyzing the fixed points and their stability we can find that there are three types of dynamical states:

1. Stable single fixed point (either low or high).
2. Two stable, one unstable fixed points (bistability).
3. Oscillations.

We can find two bifurcations of the system, where it transitions between states. The Saddle-node bifurcation indicates a transition from one stable fixed point to two stable fixed points and one unstable fixed point in-between. Its location can be identified by the points where det(Jac) = 0. The Hopf bifurcation is a transition from 1 stable fixed point to a limit cycle and is indicated by tr(Jac) = 0. Note that as long as *τ* ≪ *τ*_*w*_ the location of the bifurcations is not strongly affected by the timescales of the system. In practice, we use numerical bifurcation continuation using XPPauto [61] to obtain the bifurcation diagrams. Fig. 2a shows the bifurcation diagram for *θ* and *b*.

### E. Analytical description of the invariance in a piece-wise linear equivalent model

To simplify the analysis, we will substitute the sigmoid non-linearity with an equivalent piece-wise nonlinear function:

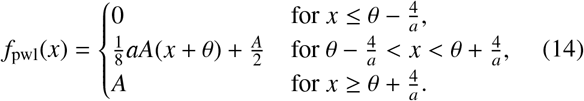

For larger values of *a*, which controls the steepness of the sigmoid function, this approximation becomes more accurate. The nullclines are now given by

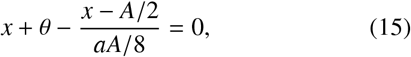

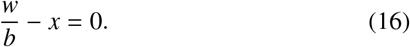

We can also explicitly solve for the fixed points,

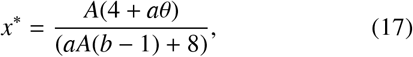

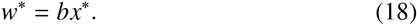

We can find the low and high values of the adaptation (*w*^−^ and *w*^+^) using the discontinuity points of the non-linearity. The adaptation variable would then be close to *w*^−^ between bursts and close to *w*^+^ during the bursts when the adaptation timescale is much slower than the activity timescale. To do that we solve the nullcline equation *w* = −*x* + *f*_pwl_(*x*). The lowest and highest points of the rate (*x*^−^ and *x*^+^ respectively) are given by 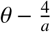 and 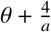. The values of the adaptation are then

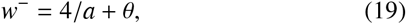

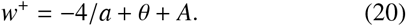

#### Invariance in the deterministic case

Next, we analyze the invariances in the model parameters using the lowest and highest values of the adaptation variable. We assume that the duration of up and down states are fixed and do not change (e.g. like in the case of oscillatory dynamics without noise cv(*T*_down_) = 0). Thus, we assume that the system jumps between two branches of the nullcline and the adaptation slowly relaxes to *w*^−^ and *w*^+^. Note that this is also a very good approximation of the original model when *a* is sufficiently large. We then analyze only the slow variable, assuming that the input 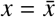 thermalizes fast and it is thus fixed and equal to 0 in the down-state and *A* in the up-state. The general solution of the initial value problem (IVP) for *w*(*t*) is

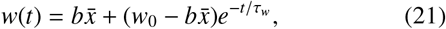

where *w*_0_ is the initial condition. Now we can solve for both *b* and *τ*_*w*_ as a function of *θ*, given fixed *T*_down_ and *T*_up_.

The time it takes to relax to *w*^−^ after the jump from the upper branch is given by the IBI, therefore *w*(*t* = *T*_down_) = *w*^−^. Thus, we can now explicitly solve for *τ*_*w*_

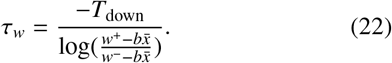

During the down state 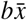 is approximately zero and given a fixed *T*_down_ the *τ*_*w*_ therefore depends only on *w*^−^ and *w*^+^, which are functions of *θ* as defined above. Thus, *τ*_*w*_ is proportional to the duration of the down state (see Fig. 9b). We can further simplify the equation and write it as

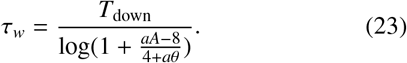

Now we can perturb *log*(1 + *u*), where 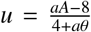 up to the first order and arrive at the equation that defines the dependency between *θ* and *τ*_*w*_

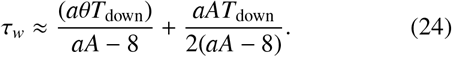

We apply a similar approach and fix the burst duration as *T*_up_. The input is then 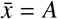 and that *w*(*t* = *T*_up_) = *w*^+^. Eq. 21 becomes

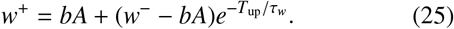

We substitute *τ*_*w*_ with Eq. 23 and get a curve of the dependency between *b* and *θ* given fixed *T*_up_ and *T*_down_ (plotted in Fig. 9a)

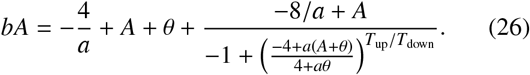

We asymptotically expand this equation up to the zero order with the Laurent series and obtain

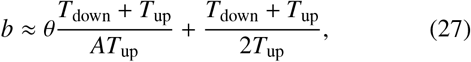

from which we can approximate *θ* as

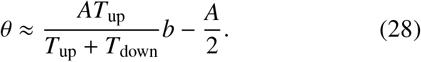

Thus, the slope of the dependency between the excitability and adaptation is 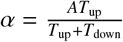.

## II. EXPERIMENTAL METHODS

### MEA recording of networks of mouse cortical neurons

The neurons of the mouse cortex (E18) were dissected and cultured according to previously published protocols [62]. The local Animal Care and Use Committee approved animal protocols for primary cell cultures. Cortical cultures were seeded on 24-well MEA plates (Multi Channel Systems MCS GmbH, Reutlingen, Germany) at the seeding density of around 5000 neurons/mm^2^. Each well contained 12 gold electrodes. The medium was changed every 3 days.

The spontaneous network activity was recorded at 17 and 18 DIV in the recording medium containing 140mM NaCl, 4.2mM KCl, 2mM CaCl_2_*2H_2_O, 1mM MgSO_3_*7H_2_O, 0.5mM Na_2_ HPO_4_, 0.45mM, NaH_2_PO_4_, 5mM HEPES and 10mM Glucose (pH was controlled to be within 7.35-7.45 range by adding NaOH). After the MEAs were transferred to the amplifier, we let the network stabilize for 10 minutes. The recordings were done while keeping the temperature at 37°C. The raw signal was recorded for 20 minutes at a sampling rate of 10 kHz and was filtered by a fourth-order lowpass Butterworth filter with a cutoff frequency of 3500 Hz, and a second-order high-pass Butterworth filter with a cutoff frequency of 100 Hz. Spike detection was performed via the noise threshold method provided by the Multi-well Analyzer Software (Multi Channel Systems MCS GmbH, Reutlingen, Germany). For each recording, the default algorithm was set to calculate the standard deviation from 50 different 100 ms long segments of raw data, and the threshold for spike detection was set at ±5 times the standard deviation level. For a subset of wells, we also blocked the inhibitory activity by adding 40*μ*M of bicuculline, this data was not included in the analysis.

### Ca-recording of networks of primary rat hippocampal neurons under changing concentrations of KCl

#### Primary cultures preparation

All procedures were approved by the Weizmann Institutes Animal Care and Use Committee. The dissections and cell dissociations were done according to established protocols [63, 64]. Hippocampal neurons were obtained from Winstar rat embryos at E19.

Brains from embryos were dissected on ice in L-15 medium with 0.6mg/ml D-glucose and 20 *μ*g/ml gentamycin. Hippocampi were dissociated in papain solution (papain 100 units, DNAse 1000 units, L-Cystein 2 mg, NaOH 1M 15 *μ*L, EDTA 50 mM 100 *μ*L, CaCl_2_ 100 mM 10 *μ*L, dissection solution 10 mL) at 37*C*. After 20 minutes of incubation at 37*C* with papain, the solution was replaced with 10 mL of plating medium (MEM without glutamine supplemented with 0.6% glucose, 1% GlutaMAX, 5% Horse Serum, 5% Fetal Calf Serum and 0.1% B27) with 25 mg of trypsin inhibitor and 25 mg of Bovine Serum Albumin, that effectively stops tissue dissection. The resulting tissues were titrated with glass pipettes after which the cells were counted with an automatic cell counter.

The suspension was then seeded with a density of around 5000 neurons/mm^2^ on glass coverslips coated with poly-L-lysine placed in 24-well plates. The plates were incubated for half an hour in a humidified, 37°C and 5% CO_2_ incubator to allow the attachment of neurons. Finally, 2 mL of serumfree medium (Neurobasal, B27 4%, GlutaMAX 1%, and FCS 1%) were added and the cultures were placed back into the incubator. At 4 DIV the glial proliferation was stopped by adding in the medium 20 *μ*g*/*mL 5-fluoro-2-deoxyuridine and 50 *μ*g/mL uridine (Sigma, Israel). Cultures were fed every day, replacing 0.5 mL of the old medium with a new feeding medium containing 90% MEM and 10% of inactivated Horse serum.

#### Ca-imaging

Recordings were done between 14 and 28 DIV. The recordings were done in an external medium (EM), containing 130mM NaCl,4mM KCl, 2mM CaCl_2_, 1mM MgCl_2_, 10mM HEPES, 10mM Glucose, 45mM Surcose. On the recording day, a coverslip with culture was transferred to a petri dish with 1mM of the EM with 4*μ*L Fluo-4. The culture was then incubated in darkness for 1 hour, after which the medium was changed to 2ml of EM. Before imaging the Perti dish was mounted in Zeiss Axiovert 135TV. The images were taken every 0.0546 s. To analyze population activity, we averaged the fluorescence from the whole field of view (which typically contained about 30-200 neurons).

### KCl changes

In addition to standard EM (which contains KCl 4mM), we prepared a set of EM with modified concentrations of KCl: 1, 2.5, 5.5, 7, and 10mM. After the initial 10-minute recording with the baseline concentration, we changed the medium to a medium with a modified concentration. We let the networks adjust for 10 minutes and then recorded another 10 minutes of spontaneous activity in the new concentration of KCl. After that, we changed the medium again, back to the baseline condition. We typically start with a baseline of 4mM, but for a small number of validation experiments, we started with higher or lower concentrations. For the final averages, these data were pooled together.

## III. DATA ANALYSIS

### A. Ca-recording

To analyze the population bursts recorded with Ca-imaging, the fluorescent imaging data were averaged over the whole recording field to obtain a time series. We validated the recordings manually and excluded those that did not show population bursts in the control conditions. Then we removed the slow fluctuations of the baseline by fitting a polynomial of the 4th degree to the data and verifying the prediction of the fit from the raw data. We smooth the time series by applying a low-pass filter (digital Butterworth filter of the third order with 1Hz cut-off frequency).

Burst onsets were detected where the derivative of the detrended signal crosses a threshold of 0.5 a.u. for at least two consequent points. We then check if the maximum amplitude within the burst exceeds 3 std of the burst baseline (calculated during 1s before the identified burst onset) and discard the events if it does not. The end of the burst for Ca-data is computed as the decay of the trace to 25% of its maximum, similarly to [24]. The burst detection and duration was then validated by visual inspection.

The burst summary statistics were computed from the extracted bursts. Note that for computing the average inter-burst interval, the duration is taken from the end of one burst to the onset of the next one.

### B. MEA data

#### Datasets

Additionally to our MEA recording at 17 and 18 DIV, we analyzed several open datasets of MEA recordings for primary cell cultures [25] as well as hPSC-derived neuronal cultures [21, 42], that include both embryonic PSC and induced PSC. The data for mice cortical and hippocampal primary cell cultures were accessed from http://github.com/sje30/g2chvc. Recordings of the activity of hiPSC-derived and rat cortical neurons were downloaded from https://gin.g-node.org/NeuroGroup_TUNI/Comparative_MEA_dataset/. We used only the data recorded across development from 3 to 66 DIV: hPSC_MEA1,hPSC_MEA2,Rat_MEA1. We used the spike times detected by the authors as described in the original publications [25, 42].

#### Burst detection

We first exclude the recordings that did not have a clear bimodal distribution of the activity. To this end, we check if the bimodality coefficient [65] of the spike count across all channels in 200ms bins is above 0.4.

Next, we identify the bursts by combining the spike times from all channels and detect the events based on the minimum inter-spike interval, burst duration, number of spikes inside a burst, and the minimum inter-burst interval. The algorithm that we use is inspired by the minimum interval algorithm (MI), which has been shown to robustly detect burst [56, 66]. We first analyze all inter-spike intervals and detect the times when the inter-spike interval is below the threshold of the mean inter-spike interval for the recording. Then we combine all bursts that are closer than a minimal IBI, after which we discard all the events that are shorter than the minimum burst duration.

The inter-spike interval threshold was set for each recording to the mean of the inter-spike across all channels (but not below 50ms or greater than 500ms). The minimum number of spikes inside a burst was 45, the minimum burst duration was 50 ms, minimum inter-burst interval was 500 ms. For the hPSC dataset, [42] we adjusted the minimum number of spikes in a burst to 100ms and the minimum possible interspike interval threshold to 20ms which allowed us to separate longer better bursts. The detection for all datasets was validated by visual inspection.

## IV. EXTENDED FIGURES

**FIG. 7.**
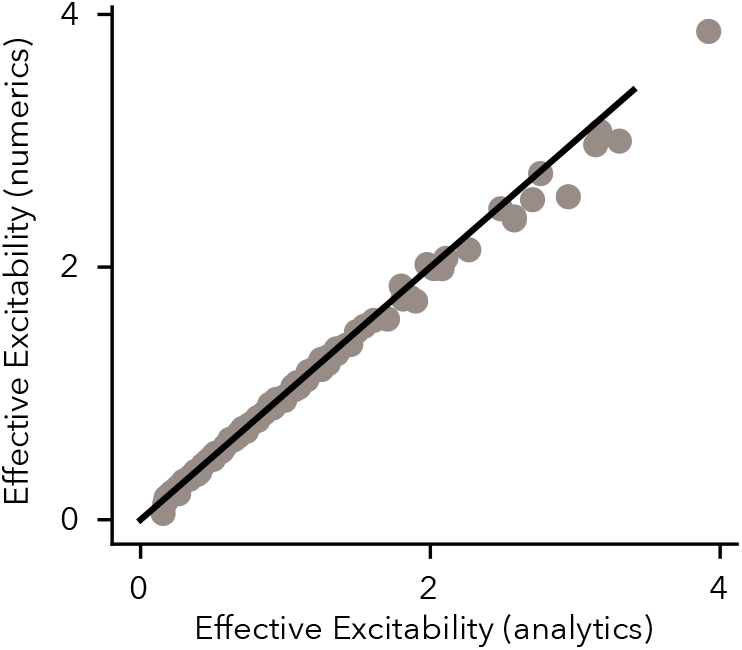
Effective excitability estimated from the average summary statistics analytically closely follows the slope of the dependency between excitability and adaptation strength (numerics) estimated from the posterior samples. In this plot we uniformly sampled summary statistics (IBI between 3 and 20s, burst duration between 1 and 3s, and CV of IBI between 0,1) and then sampled from the corresponding posterior distributions. The numerical effective excitability was estimated as the slope of the best-fitting line for the *θ* and *b* samples. Analytical values were computed as *AT*_up_*/*(*T*_down_ + *T*_up_).

**FIG. 8.**
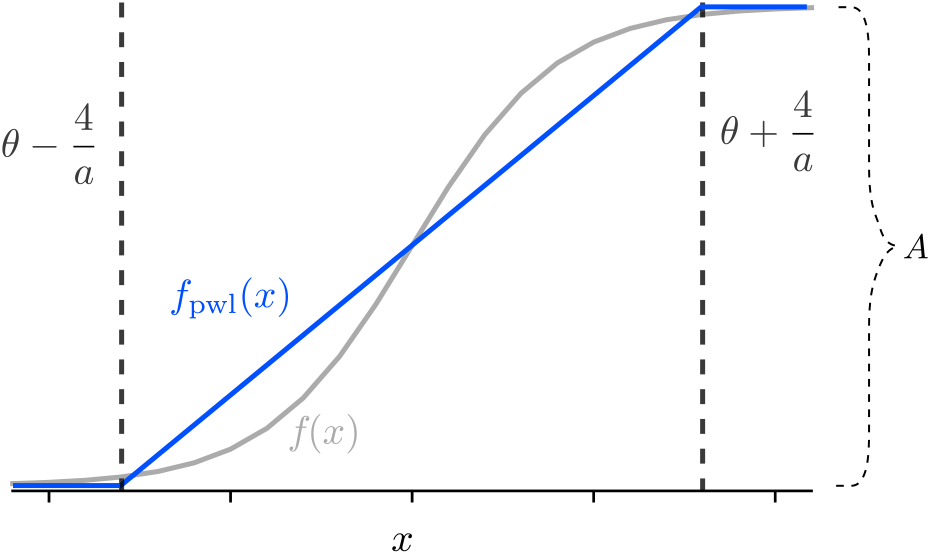
Piecewise linear approximation ( *f*_pwl_(*x*)) of the original sigmoid ( *f* (*x*)).

**FIG. 9.**
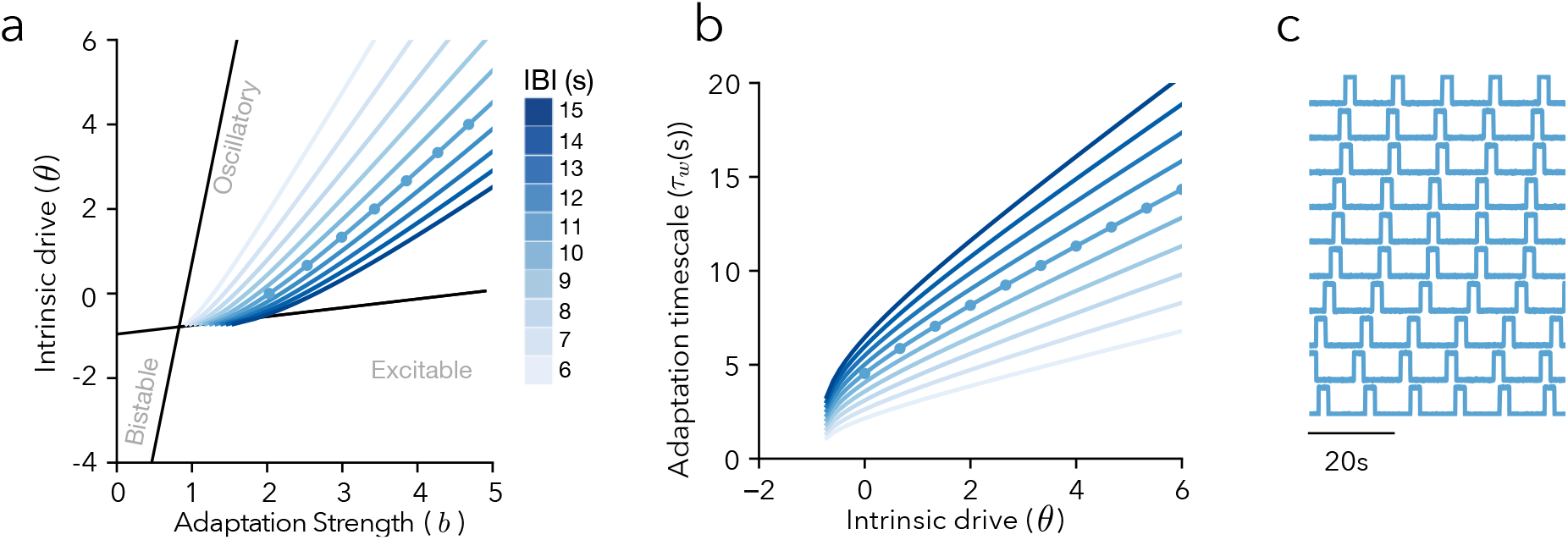
The invariance of bursting activity in an oscillatory regime without noise. **a**) Solution of the adaptation strength of a function of *θ* for different fixed values of IBI and 2.5s burst durations **b**) Solutions for the *τ*_w_ as a function of *θ*. **c**) Examples of traces with the IBI=10s and low noise (*σ* = 0.1)

**FIG. 10.**
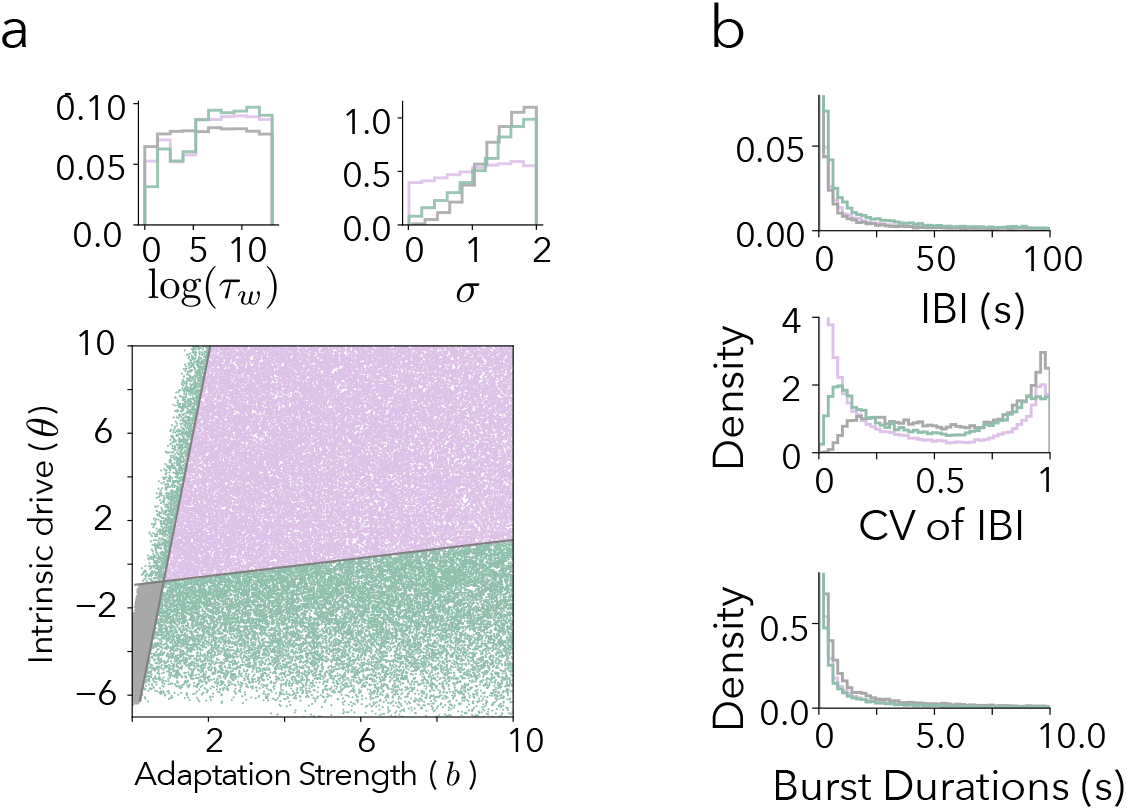
The simplified model can produce a wide range of network bursting statistics.**a** Samples from the prior of the model parameters that lead to bursting activity (pink - oscillatory state, green - excitable state, gray - bistability). **b** Histogram of corresponding summary statistics.

**FIG. 11.**
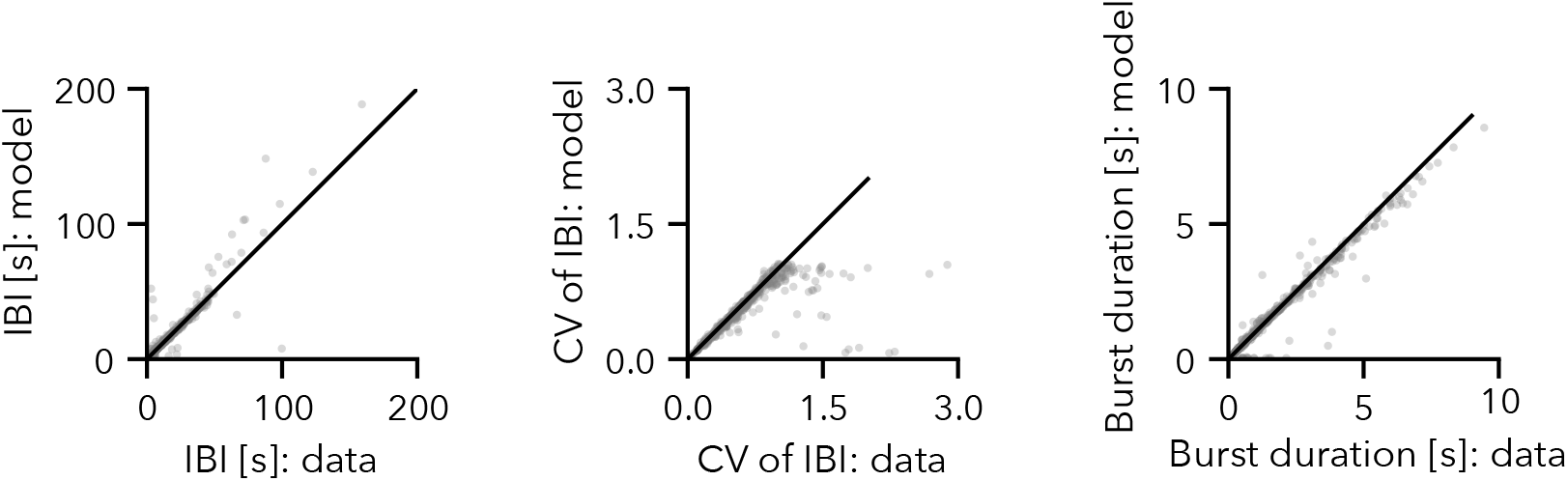
Performance of the maximum aposteriori values for the posteriors fitted to the cortical, hippocampal, hPSC cultures’ activity from Fig. 3.

**FIG. 12.**
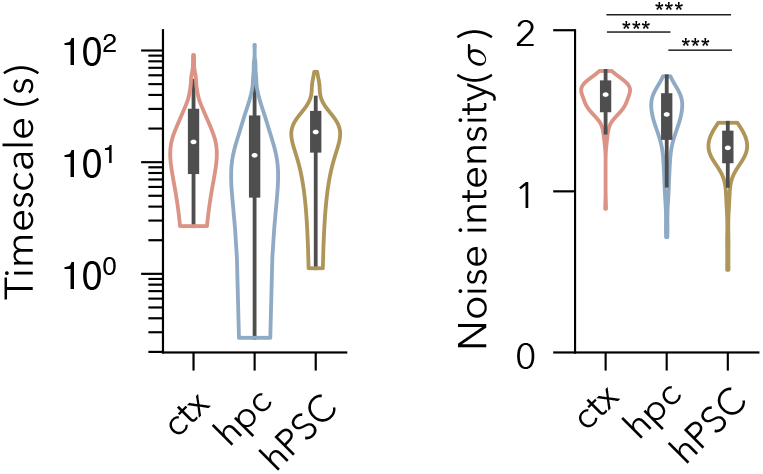
Timescale and the noise intensity for the inferred model parameters for cultures of mouse primary neurons from cortex, hippocampus (>14DIV)[25] and hPSC (>31DIV) [42] of Fig. 3. Here, we extract the timescales by computing the slope of the dependency between *b* and *τ*_*w*_ from the fitted posteriors. We could not reject the null hypothesis of the mean equality between the timescales of cultures types (One-way ANOVA F(2,375) = 2.206, p=0.11). The noise intensities were significantly different between all groups (One-way ANOVA F(2,375) = 115.6, *p <* 0.00001, pairwise ind. t-test ctx vs hpc t-val = 8.8, p *<* 0.00001; ctx vs hiPSC t-val = 9.158 p < 0.00001; hpc vs hiPSC t-val=8, p < 0.00001.)

**FIG. 13.**
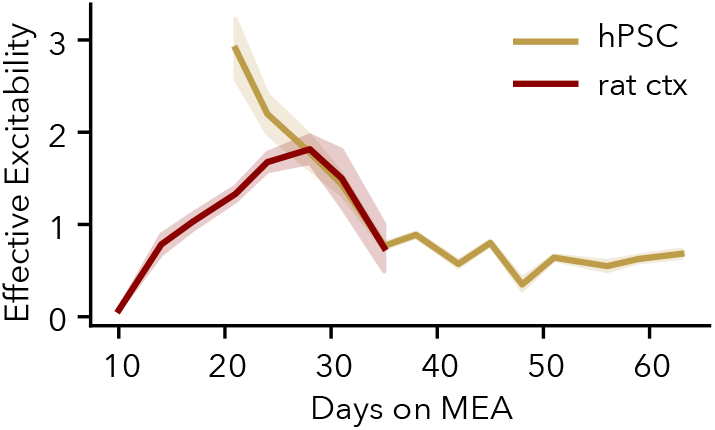
Developmental profiles of the population bursting development in hPSC and primary *in vitro* networks of rat primary cortical neurons from the same dataset [21, 42]. hPSC are the same as in Fig. 4 (shaded area - s.e.m.)

**FIG. 14.**
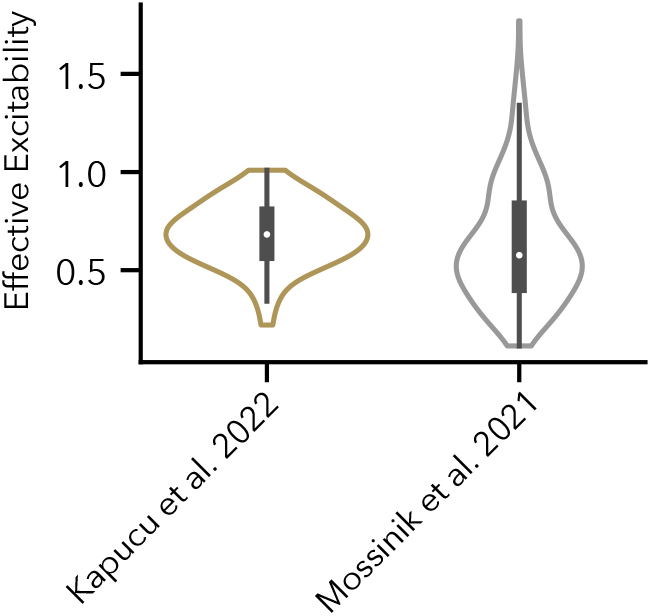
Comparison of the effective excitability between hPSC datasets. induced hPSC from [22] (21 DIV, N=286) and induced and embryonic hPSC [21, 42] (cultures older than 31 DIV, N=55). Two-sided ind. t-test t=1.34, p=0.18 (not corrected).

**FIG. 15.**
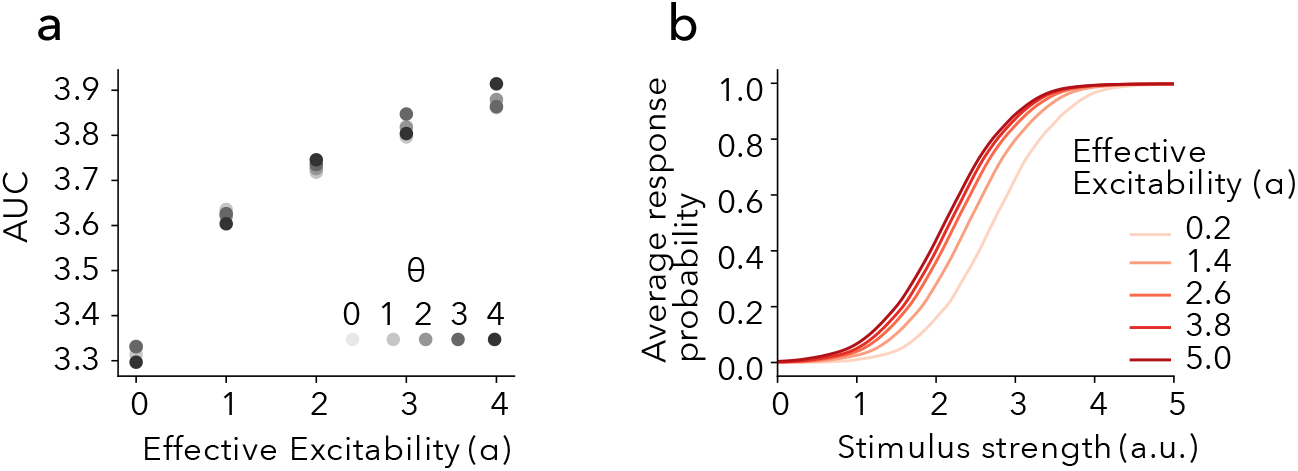
The effective excitability sets the response properties of neuronal networks. **a)** Area under the curve (AUC) of responses (computed with trapezoid rule) for increasing effective excitability and different levels of the intrinsic drive (*θ*). **b)** Leftward shift in the response probability in models with increasing effective excitability(*α*).

**FIG. 16.**
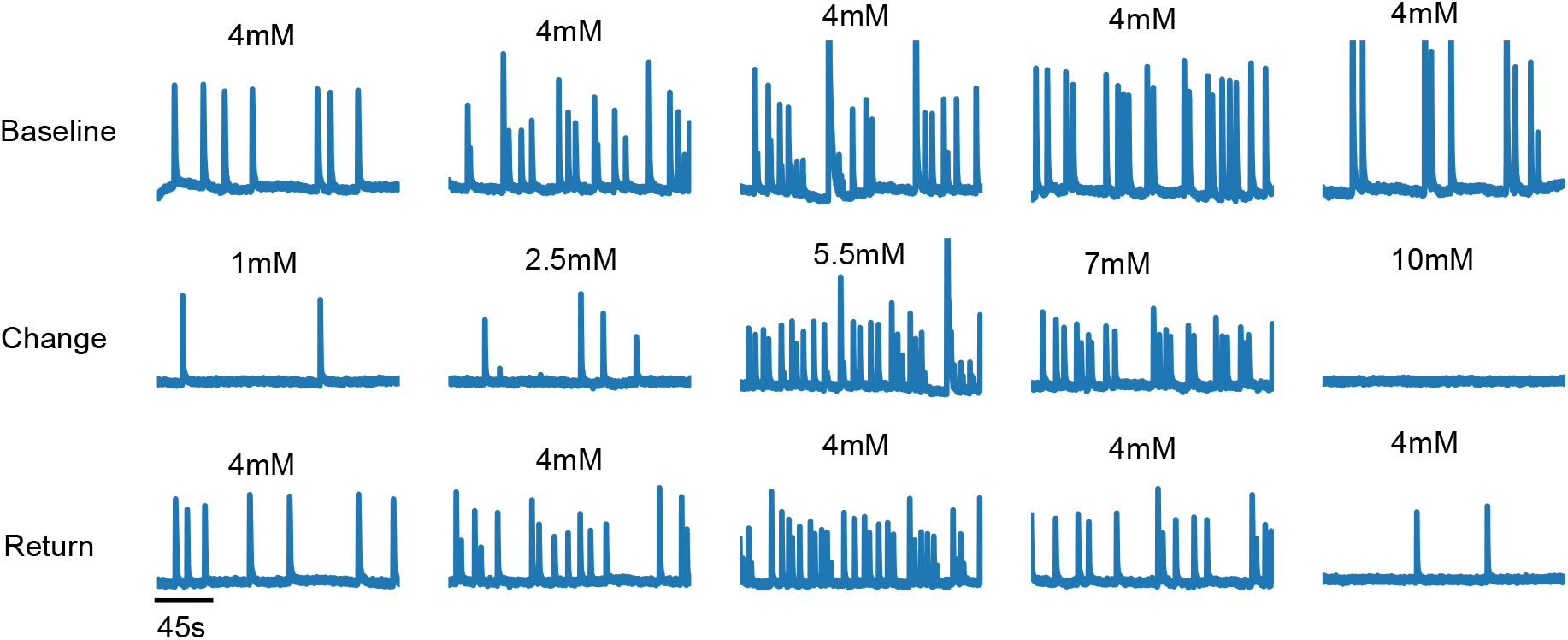
Examples of average Ca-traces in experiments with different concentrations of KCl and return to the baseline condition.

